# General prediction of T cell receptor antigen specificity from sequence using AlphaFold 3

**DOI:** 10.64898/2026.06.02.729478

**Authors:** Lawson J. Woods, Brandon Neff, Kamel Lahouel, Antoine Goursaud, Mete Mulazimoglu, Kameron Bates, Cristian Tomasetti, John A. Altin

## Abstract

The Major Histocompatibility Complex (MHC):peptide:T Cell Receptor (TCR) complex is the most diverse trimolecular interface known in nature and a key trigger for adaptive immunity. TCRs are now routinely sequenced at scale, however decoding their specificity for antigens remains a bottleneck. While *in silico* approaches have advanced considerably, none enables prediction against “unseen” epitopes, limiting their applicability to a tiny fraction of cases. Here, we show that AlphaFold 3 (AF3) predicts the structures of MHC:peptide:TCR triads with unprecedented accuracy. By applying AF3 to >9,000 TCRs mapped to >1,000 distinct epitopes restricted by >70 MHC class I / II alleles, we identify features that distinguish cognate triads from controls. The resulting model achieves median AUCs of 0.81-0.92 on validation triads unseen by AF3 or during feature selection. Our results reveal that generalized prediction of TCR specificity from sequence is possible, with the potential to greatly accelerate the decoding of immune responses.

## Introduction

Adaptive immunity relies on lymphocyte antigen receptor repertoires that are broad enough to address a vast array of potential pathogens, yet also precise enough to distinguish between closely-related antigens^1,2^. In the case of T cells, this feat of highly-sensitive and specific sensing is accomplished within each individual by >10^8^ unique T Cell Receptor (TCR)α:β heterodimeric proteins (among an estimated total of >10^15^ possibilities) that can collectively recognize and distinguish among billions of peptide antigens presented by diverse Major Histocompatibility Complex (MHC) proteins^3,4^. The study of TCR specificity for p:MHC antigen is an important focus in basic immunology^5^, and forms the basis for powerful new diagnostic^6^,^7^ and therapeutic^8^,^9^ approaches.

However, the vast diversity of the MHC:peptide:TCR system poses several challenges. The first is that TCRs of interest are typically admixed within complex repertoires. Recent advances in high capacity sequencing and single cell genomics have allowed TCR repertoires to be routinely sequenced, with increasing depth, and increasingly in a way that preserves the pairing of TCRα and β chains^10^. However, a second challenge is to translate the resulting sequences into antigen specificities, which has emerged as a key bottleneck^11^. Experimental methods for linking TCRs to antigen specificity include binding assays that use p:MHC probes^12,13^, as well as cell culture systems that use proteins or peptides to activate reporters in an antigen-specific fashion^14^. Although recent embodiments have leveraged DNA barcoding to multiplex antigenic diversity to high dimensionality^12,15,16^, even the most sophisticated experimental approaches typically take months to query only a small number of TCRs.

In response to these challenges, the past decade has also seen parallel and increasingly fruitful efforts to develop purely computational approaches to solve the TCR ‘sequence to specificity’ problem^17^. These methods generally fall into two categories. The first category includes algorithms such as TCRdist^18^, GLIPH^19^, DeepTCR^20^, TCRAI^21^, among several others^22–26^, that identify TCR sequence features associated with specificity for particular p:MHC antigens, and use these features to organize newly-observed TCRs into groups to make inferences about their specificity. Although powerful, the utility of these methods for specificity prediction is limited in two important ways: first, they do not generalize across antigens and so require that a model be pre-trained on known TCRs for each antigen of interest, and second, they are often insensitive to subsets of TCRs whose sequence features do not form detectable clusters yet are antigen-specific^16^.

The second category of methods for predicting TCR specificity from sequence uses 3-dimensional molecular modeling to estimate the degree of structural complementarity between a TCR and a set of candidate p:MHC antigens. Insofar as they take account of protein structure explicitly, these approaches are conceptually advantageous over non-structural methods. However, while some models require experimental or templated structural data[27, 28], more general approaches rely on accurate prediction of the 3-dimensional folding and docking of MHC:peptide:TCR interfaces, which has been a challenging problem to solve. Earlier approaches showed significant, albeit often modest, performance on well-characterized epitopes^29,30^. More recently, powerful advances in accurate protein structural prediction driven by deep neural networks, especially AlphaFold and its progeny, have been adapted to the study of MHC:peptide:TCR complexes, with promising results^31–35^. However, even here, demonstrated successes on the specificity prediction task have been restricted to a small number of well-studied and structurally characterized p:MHC epitopes. This may be attributable in part to the fact that even the most advanced of the implemented models (AlphaFold-Multimer^36^) relies heavily on residue covariation, which is not well-represented during training or inference because the available sequence alignments and 3-dimensional structures sample only a tiny fraction of the diversity that is possible at the MHC:peptide:TCR interface^17^. Therefore, despite important progress, the prediction of TCR specificity for “unseen epitopes” from sequence alone remains an unsolved “grand challenge” in immunology^37^.

AlphaFold 3 (AF3) is a recently developed model that improves on earlier versions in part by simplifying the way aligned sequences are represented and introducing a generative diffusion step to refine the predicted atomic coordinates^38^. AF3 shows less reliance on sequence alignment and far superior performance on protein complexes, including antibody-antigen complexes, properties that we reasoned could also enable general prediction of TCR specificity. Here, we test this hypothesis by applying AF3 to large and diverse sets of complexes (“triads”) formed by MHCI/II, peptide and TCR. We use structurally-characterized triads from PDB (Protein Data Bank) to benchmark structural prediction accuracy, and then a much larger and more diverse set of less characterized triads from IEDB to identify features that correlate with TCR status (cognate v non-cognate). We characterize heterogeneity within the IEDB, and finally benchmark performance of the best classification model on separate hold-out sets of class I- and II-restricted triads from high-quality assays.

## Results

### Benchmarking AF3 performance on MHC:peptide:TCR triads with known structures

To test the performance of AF3 in accurately modeling the structures of MHC:peptide:TCR complexes, we began by analyzing solved MHC:peptide:TCR structures from the PDB. We used the TCR3d database [39] to comprehensively identify 284 solved ternary structures: 214 MHC class I and 70 MHC class II. Of these, a subset of 129 (89 MHC class I and 40 MHC class II) had been previously processed by the AlphaFold-Multimer structural prediction pipeline specialized for MHC:peptide:TCR complexes (AF-TCRdock)^31^. PDB codes, MHC, peptide and TCR sequences for all solved triads that were analyzed are provided in **Supplementary Table 1**.

For each triad, we used AF3 to predict the structure (from sequence) of the protein complex formed by its 5 constituent protein chains: peptide, MHCα, MHCβ, TCRα, TCRβ. We found that the structures output by AF3 consistently resembled the expected MHC:peptide:TCR ternary complex at a broad spatial scale, with each predicted structure characterized by: (i) the expected overall folding and pairing of chains to form the individual TCRα:β and MHCα:β heterodimers, (ii) localization of the peptide within the binding groove of the MHC, and (iii) the formation of an interface between the MHC alpha helices, the peptide, and the TCR Complementarity Determining Region (CDR) loops (**Figure 1a**).

**Figure 1:**
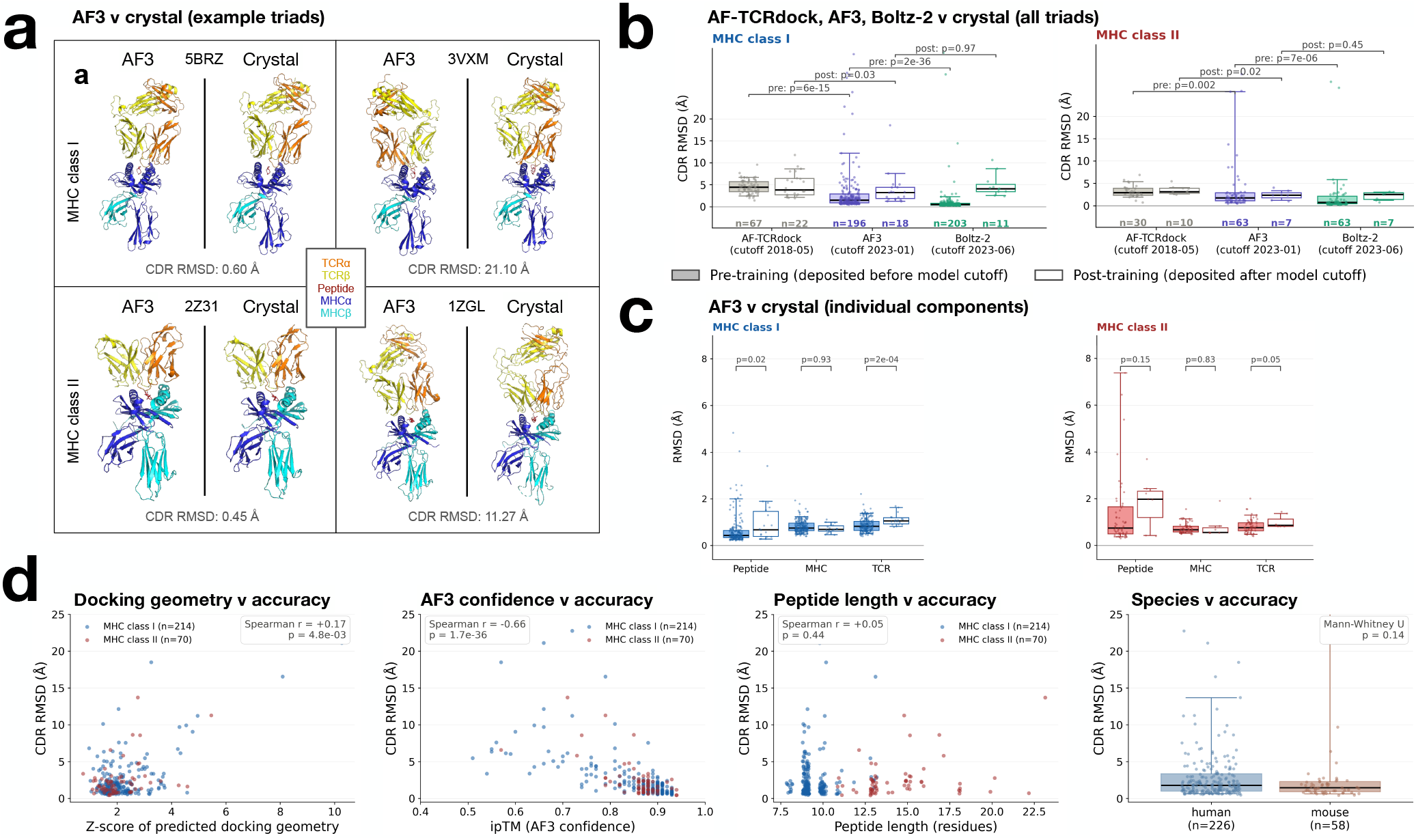
AF3 predicts MHC:peptide:TCR structures with unprecedented accuracy that is reliably estimated by confidence metrics. **a**, Example MHC:peptide:TCR triad structures predicted by AlphaFold3 (*left* in each pair), and their known crystal structure counterparts (*right* in each pair). For each MHC class (*rows*), shown are the triads with the most (*left*) and least (*right*) accurate AF3-predicted structures measured by the CDR RMSD method that compares the arrangement of the TCR’s hypervariable loops relative to the co-ordinate frame of the peptide:MHC antigen^31^. Note that 3VXM’s predicted structure is reverse-docked with respect to the canonical orientation present in the true structure. **b**, Comparison of the performances of AlphaFold-Multimer (TCRdock), AlphaFold 3, and Boltz-2, comparing predicted v crystal structures using the CDR RMSD metric. Triads predicted with each method are categorized according to when they were deposited in PDB relative to the training of the respective model. For the TCRdock RMSDs, we used values calculated in a previous study^31^. Annotated p-values are generated using the Wilcoxon rank-sum test. **c**, RMSD analysis of all 284 triads analyzed by AF3 (214 MHC class I, 70 MHC class II), comparing predicted structures and crystal structures, now analyzed at the level of individual protein chains. MHC and TCR values are based on alignments of those components individually, whereas peptide values are calculated based on prior alignment of the peptide to the MHC’s co-ordinate frame (rare outliers result from the prediction of incorrect peptide binding registers). **d**, Correlates of AF3 performance, evaluated over the 284 triads described in (c) and measured against the CDR RMSD metric (common y-axes). We observe a strong correlation with overall AF3-predicted structural confidence (captured by the ipTM summary metric), and a weaker but still significant correlation with Z-score of docking geometry (calculated as described in the **Methods**) (*left 2 plots*). No correlation is observed for peptide length or triad species of origin (*right 2 plots*).

To quantify the accuracy of the AF3-predicted structures and benchmark their quality in comparison to other methods, we first performed Root Mean Square Deviation (RMSD) analysis comparing the predicted structures with the corresponding crystal structures in PDB, using a previously-described method that focuses on the arrangement of the TCR’s Complementarity Determining Regions (CDRs) with respect to the p:MHC complex (CDR RMSD)^31^. We compared the performance of AF3 against the current state-of-the-art model (AF-TCRdock, which consists of AF2-multimer with curated chain templates^31^), as well as a recently-released alternative model to AF3 with broadly similar overall performance (Boltz-2^40^). We studied the generality of each model beyond its training data by partitioning the triads into 2 model-specific groups: those released before the model’s training cutoff date (therefore included in its training), and those released afterwards (therefore not included).

Overall, AF3 performed very well, yielding median predicted v crystal CDR RMSDs of 1.5Å and 1.8Å for the pre-training triads, and 3.2Å and 2.4Å for the post-training triads (for classes I and II, respectively) (**Figure 1b**). In all four cases, AF3 significantly outperformed AF-TCRdock, with median RMSD improvements of 0.8Å – 2.9Å (6e-15 < p < 0.03). While no decrease in performance on triads from post-v pre-AF3 training cutoff was observed for AF-TCRdock in either MHC class I or II (p=0.51 for class I, p=0.21 for class II), AF3 showed slightly reduced performance on MHC class I post-cutoff triads (class I p=1.6e-3, class II p=0.26), perhaps because AF-TCRdock performs inference with curated template information that informs structure predictions. Boltz-2 showed decreased performance on post-training triads relative to pre-training triads for both classes (class I p=1e-7, class II p=0.01). While it strongly outperformed AF3 on pre-training triads (p=2e-36 and 7e-6), there was no improvement on post-training triads (p=0.97 and 0.45), suggesting that AF3 has achieved a greater degree of generalization from its training data.

We next analyzed AF3’s prediction accuracy for individual components of the triad: the TCR and MHC chains aligned individually, as well as the peptide individually after alignment on the MHC (**Figure 1c** – see **Methods** for alignment details). For both class I and class II structures, concordance between known and predicted structures of these individual elements was excellent, with median RMSD values ranging from 0.4-1.0Å for all components and classes (with the exception of peptides of post-training class II triads where RMSD=2.0Å). Moreover, when comparing structures released pre v post the AF3 training cutoff, we observed significant (but slight) decreases in performance for the TCRs from both classes, and the peptides from class I triads. The overall higher values for interface RMSDs (CDR, peptide) compared to single-chain RMSDs (TCR, MHC) are consistent with the expectation that predicting the arrangement of subunits represents a harder challenge than predicting the structures of individual chains.

Because RMSD analysis requires a known structure and so is not available as an estimate of model accuracy in the general case, we sought to establish a relationship between AF3-predicted properties and model quality. We found that both AF3’s ipTM (interface predicted template modeling) and the Z-score of the AF3-predicted docking geometry with respect to the per-MHC-class consensus distribution extracted from the database of PDB triads in a recent study^31^ were significantly correlated with CDR RMSD as a metric for model quality (**Figure 1d**). Together, these observations indicate that features of the AF3-predicted structures can be used to assess the confidence of the prediction.

Overall, these results reveal that AF3 can predict the ternary structures of MHC:peptide:TCR complexes with unprecedented accuracy (substantially higher accuracy than AlphaFold-Multimer, even with TCRdock improvements) and frequently with precision and associated confidence outputs that are likely to be compatible with the development of a binding specificity model.

### Application of AF3 to a large IEDB dataset and identification of classification features

Although useful for initial benchmarking, the relatively small number and restricted antigen diversity of PDB structures limits their utility for the development and testing of a general model of TCR specificity. We therefore turned to a much larger dataset consisting of predominantly non-structurally characterized MHC:p:TCRs (hereafter, “cognate triads”) identified using a variety of assays across the scientific literature and curated in the Immune Epitope Database (IEDB) (^41^). We constructed a “training” dataset by querying IEDB for all cases where a human T cell response consisting of a peptide epitope restricted by a defined MHC I or II allele (resolved to field 2: a unique amino acid sequence), has been experimentally matched with reactive TCRα:β pairs (total of 8,104 class I and 1,688 class II eligible triads). To ensure generalizability, we removed triads sharing substantial sequence identity with PDB triads present prior to the AF3 training cutoff (sequence identity of ≥ 95% for MHC and TCR chains and a match ≥ 77% for peptides), resulting in 393 class I and 63 class II exclusions.

For each MHC class, we also identified separate, high-quality “validation” datasets that were left untouched to enable subsequent benchmarking of the performance of our best models. These datasets consistent of 16 class I cognate triads from PDB (deposited after the AF3 training cutoff date) and 205 class II cognate triads identified using a high-confidence single cell approach (validation datasets are described in further detail below). To maximize the stringency of these validation datasets, we also removed from the respective training datasets any triads sharing sequence identity using the same thresholds described above for the PDB triads, resulting in 8 class I training set exclusions and 6 class II training set exclusions.

As controls for both the training and validation datasets, for each unique target antigen (p:MHC), we used a permutation approach to identify a set of TCRs from the same database source (e.g. PDB, IEDB) that was *not* associated with the focal antigen (“non-cognate triads”, which consisted of 1-100-fold more triads than the cognate set for each antigen, see **Methods** for details). In total, after applying the sequence identity exclusions described, we were left with training sets of 7,702 cognate triads and 157,025 non-cognate triads across 38 MHC I alleles for class I, and 1,625 cognate triads and 33,628 non-cognate triads for class II. The validation set consisted of 16 cognate triads and 181 non-cognate triads for class I, and 205 cognate triads and 1,228 non-cognate triads across 33 MHC II alleles for class II (described further below). MHC, peptide and TCR sequences for all cognate and non-cognate triads are provided in **Supplementary Table 2** (training dataset) and **Supplementary Table 3** (validation dataset). For both the training and validation datasets, the sources and numbers of cognate and non-cognate triads, as well as strategies to avoid data leakage, are summarized in **Figure 2**.

**Figure 2:**
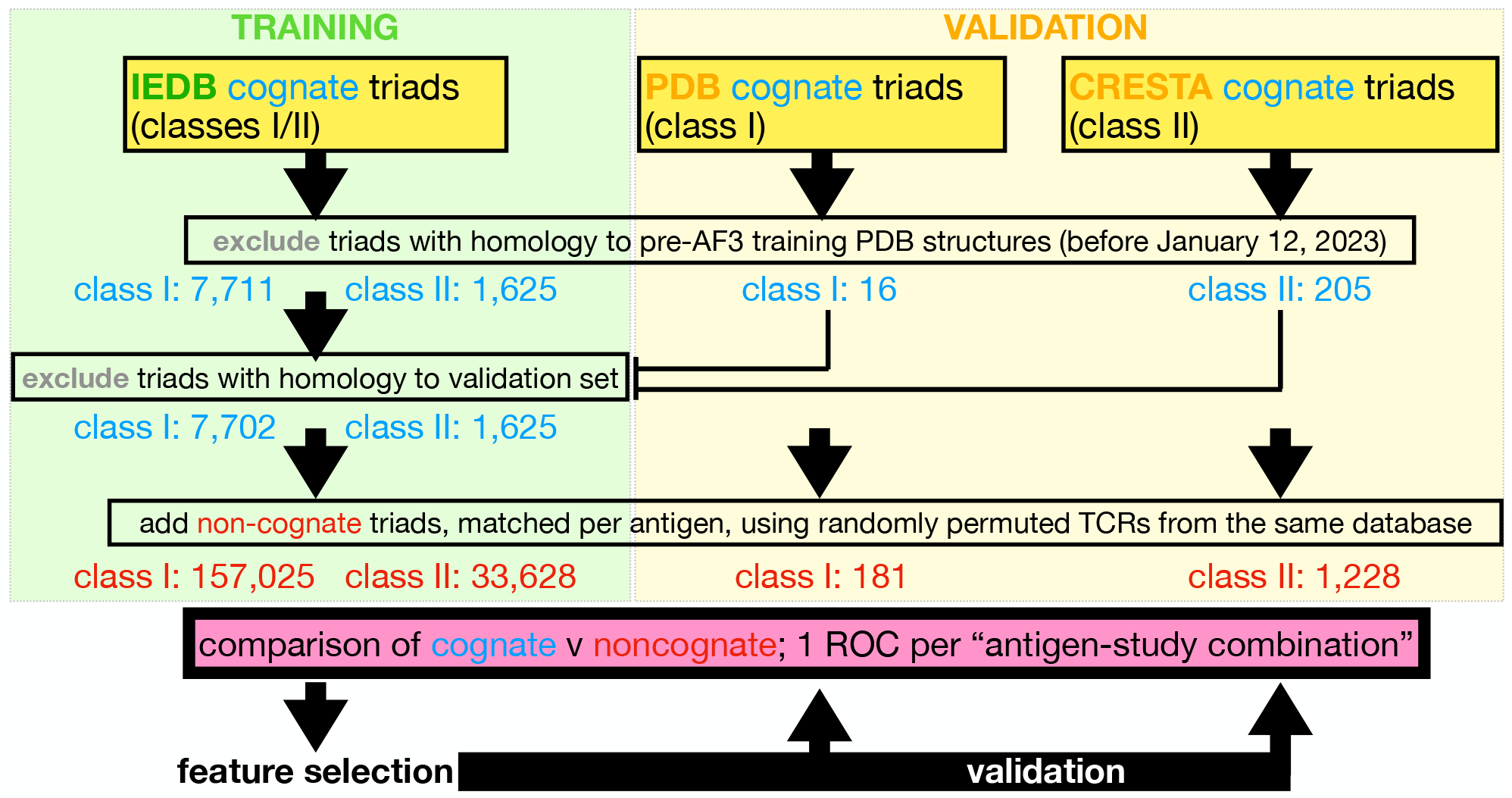
Data sources used to train and validate a general predictor of TCR specificity. Sets of cognate MHC:peptide:TCR complexes (“triads”) were sourced from IEDB (for “training”, left side) or PDB and CRESTA [16] (for “validation”, right side). To ensure generality against AF3’s training data, we excluded triads with homology to pre-AF3 training PDB structures from all training and validation datasets. To ensure generality against our training data, from the validation set we also excluded triads with homology to the training set. Within each category, we then added non-cognate controls from the corresponding dataset, in which pairings between TCRs and p:MHC antigens were randomly permuted. Data stratified by “antigen-study combination” (sets of triads identified for a particular antigen in a particular study) were evaluated by ROC analysis comparing cognate v non-cognate antigens.

To quantify the degree to which cognate triads can be distinguished from non-cognate triads, we performed AF3 predictions on all cognate and non-cognate triads from the training set and then isolated a large set of features for extraction from each prediction. These features were selected to represent aspects of both structure and the associated confidence, quantified in different regions and at different scales (**Figure 3a**). First, we used AF3’s scalar summary confidence metrics (*n* = 3), comprising: pTM (predicted template modeling), ipTM (interface pTM), and ranking score [38]. Next, focusing on the binding interface as the key determinant of specificity, we used confidence metrics associated with residues near the peptide:MHC and peptide:TCR interfaces (local metrics, *n* = 11 for triads with class I MHCs and *n* = 12 for class II MHCs) as well as confidence metrics associated with pairwise residue interactions near these interfaces (interface metrics, *n* = 26). Finally, we represented the large-scale orientation and position of the TCR relative to the p:MHC antigen using a 6-parameter docking representation (docking geometry) described previously^31^ and further extracted a Z-score of each docking geometry with respect to the consensus distribution (*n* = 7 total docking parameters). Detailed descriptions of each feature are provided in the **Methods** section.

**Figure 3:**
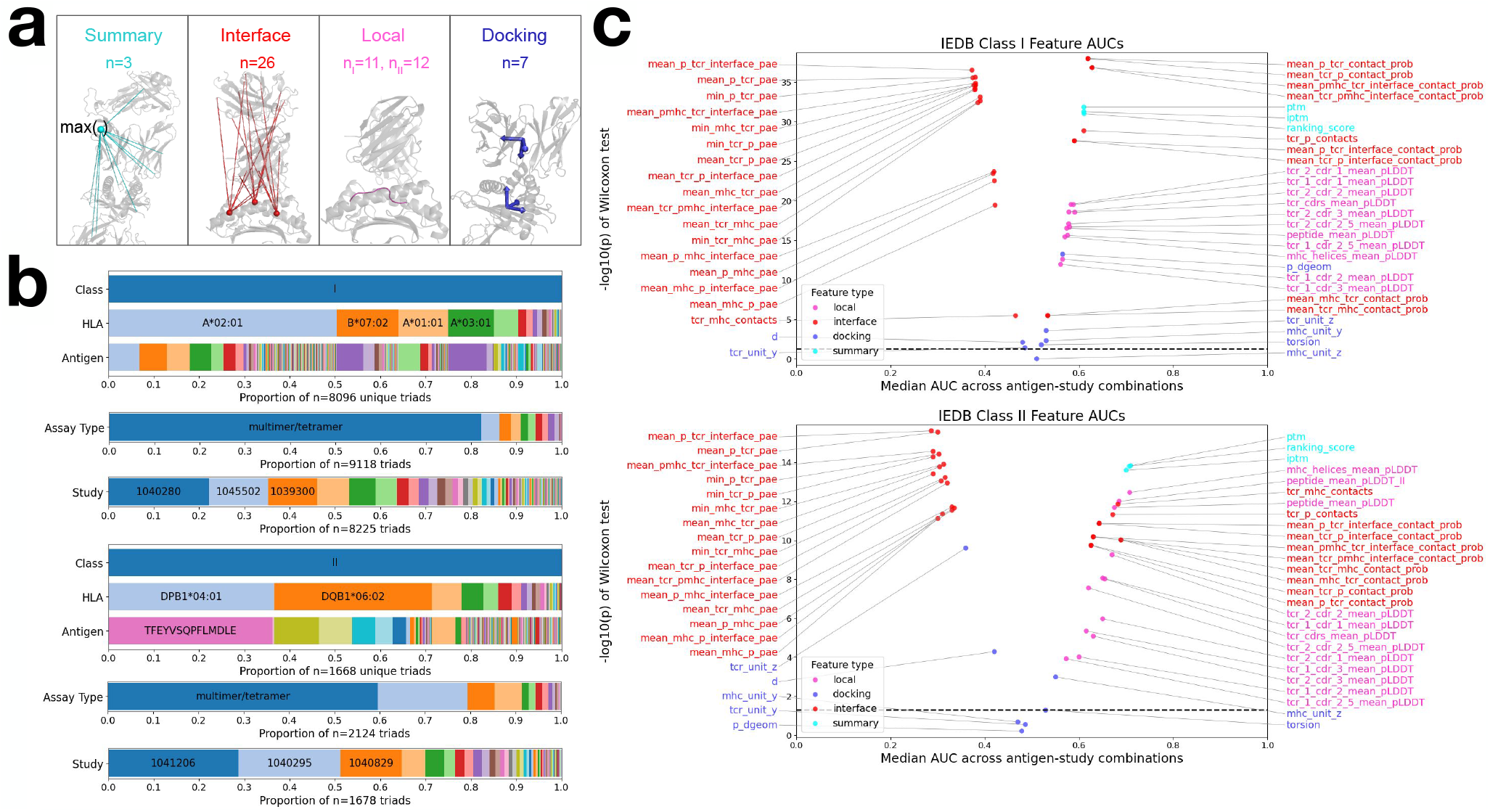
Features of the AF3-predicted MHC:peptide:TCR structure strongly correlate with cognate v non-cognate status across diverse class I-and II-restricted antigens. **a**, Overview of the four categories of features used to distinguish binding and non-binding triads. Summary features are scalar values output directly by AF3 (ipTM depicted as an example). Interface scores are curated slices of AF3 pairwise residue output arrays (mean peptide:TCR PAE depicted). Local scores are curated slices of AF3 per-residue or per-atom output arrays (mean peptide pLDDT depcited). Finally, docking scores consist of 6 geometry features per triad, generated using the TCRDock pipeline^31^. Full definitions of each feature are provided in **Methods. b**, Composition of the IEDB “training” dataset according to MHC allele, antigen, assay type and study for MHC class I (*upper*) and class II (*lower*). Note the skew towards a subset of studies and antigens (e.g. 38% of IEDB class II triads come from a single SARS-CoV-2 antigen: DPB1*04:01:TFEYVSQPFLMDLE). Prominent MHC alleles, antigens, assay types and studies are labeled on the corresponding bars. Total number of triads differ across bars because one triad can be measured with multiple assays or be measured in multiple studies. **c**, Volcano plots quantifying the degree to which each of the features (defined in (a) and colored using the same schema, n = 47 or 48) correlates with cognate v non-cognate status for class I (upper) and class II (lower) triads. To account for the skew in database composition shown in (b), we generate an individual AUC for each “antigen-study combination” (consisting of all TCRs specific for a given antigen within a particular study, matched with permuted non-cognate control TCRs). Correlation strength is estimated as the median of AUCs across all antigen-study combinations (*x-axis*), and significance as the p-value with respect to the null hypothesis (that the AUCs distribution of the given feature is symmetric around 0.5), quantified by Wilcoxon signed-rank test (*y-axis*).

IEDB represents a diverse database compiled from the scientific literature, with underlying content that was iscovered by different labs using a variety of assay types and reagents^41^. Since previous work has shown that TCR databases of this type include some content that is not reproducible^42^, we began with the assumption that a subset of the underlying literature studies, and potentially of antigens within each study, may be unreliable. Given that the distribution of cognate triads in IEDB across studies and antigens follows a highly skewed distribution (e.g. >50% of all class I and II triads come from 4/182 (2.2%) and 2/62 (3.2%) of studies, and 11/1031 (1.1%) and 3/279 (1.1%) of antigens, respectively – **Figure 3b**), treating all triads equally would bias the results heavily towards a small subset of the studies / antigens. To avoid this, we instead resolved our scoring to the level of *individual antigens in individual studies*, each of which corresponds to at least 1 cognate TCR (although frequently more). Focusing on these “antigen-study combinations” as our basic unit of analysis corresponds to the assumption that each antigen in each study has an approximately equal chance of being a reliable detector of cognate TCRs.

In total, after exclusions, the IEDB class I and II sets consisted of 1,024 and 248 antigen-study combinations, containing 1-689 (mean 7.5) and 1-378 (mean 6.5) cognate triads, respectively. For each antigen-study combination, we used Receiver Operator Curve (ROC) analysis to compare feature scores of the corresponding cognate v non-cognate triads and generate Areas Under the Curve (AUCs). Importantly, we designed our non-cognate triad selection method to ensure that each AUC was resolved to 0.01 units or better (for example, antigen-study combinations with only a single cognate triad were compared against 100 non-cognate triad controls - see **Methods**). For each of the AF3 metrics, we report the median of AUCs across all antigen-study combinations (**Figure 3c**, *x-axis*), and estimate significance by testing the null hypothesis that the AUCs for that metric come from a distribution centered about 0.5 (**Figure 3c**, *y-axis*).

Across both MHC classes, we observed significant deviations from the null hypothesis (9.9e-39 < p < 0.05) for the vast majority of features (46/47 for class I, and 44/48 for class II). These correspond to strong deviations from AUC=0.5 (as high as 0.63 for mean peptide-MHC:TCR interface contact probability in class I and as low as 0.29 for mean peptide:TCR interface PAE in class II, noting that the latter metric negatively correlates with cognate status). Strong performance was consistently observed for metrics incorporating Predicted Alignment Error (PAE) at the interface between the TCR chains and the p:MHC complex (**Figure 3c**, *red dots*). However, we also observed significant discrimination for all of the summary and local features and 6 and 3 of the 7 docking features for class I and II, respectively.

Pairwise analysis revealed strong correlations among many features, particularly the best performing ones (**Supplementary Figure 1a**. We also observed that feature ranking was stable even with downsampling to relatively small fractions of the training dataset (**Supplementary Figure 1b**). Consistent with these observations, our attempts to combine features using Random Forest and XGBoost models showed inconsistent performance benefits compared to the individual features (**Supplementary Figure 1c**). To avoid overfitting, we therefore selected the single best feature – AF3-PTI-PAE (31− *mean*_*p*_*tcr*_*interface*_*pae*, inverted using the maximum PAE bin to account for the negative correlation with cognate status) – as our best model and the basis for all subsequent classification tasks.

### Analysis of AF3 classifier performance across assay types, TCR:antigen affinities and cross-reactive peptides

Having established an AF3-based classifier (AF3-PTI-PAE) for distinguishing cognate from non-cognate MHC:peptide:TCR triads, we next used it to study heterogeneity within the IEDB dataset. To this end, we further resolved the class I and II antigen-study combinations described above according to the assay types that were used to identify their constituent triads (as reported in IEDB). In total, this consisted of 12 assay types with 4-790 antigen-study combinations each for class I, and 7 assay types with 4-89 antigen-study combinations each for class II (**Figure 4a,b**). Comparing across groups revealed that AF3-PTI-PAE classification performance was strongly dependent on assay type (p=3.2 ×10^−6^, 3.8 ×10^−14^ for class I, II), with a subset of assays showing high performance (median AUC > 0.9 in 2 cases), and others showing performance that was closer to random guess (median AUC < 0.6 for 4 class II assay types).

**Figure 4:**
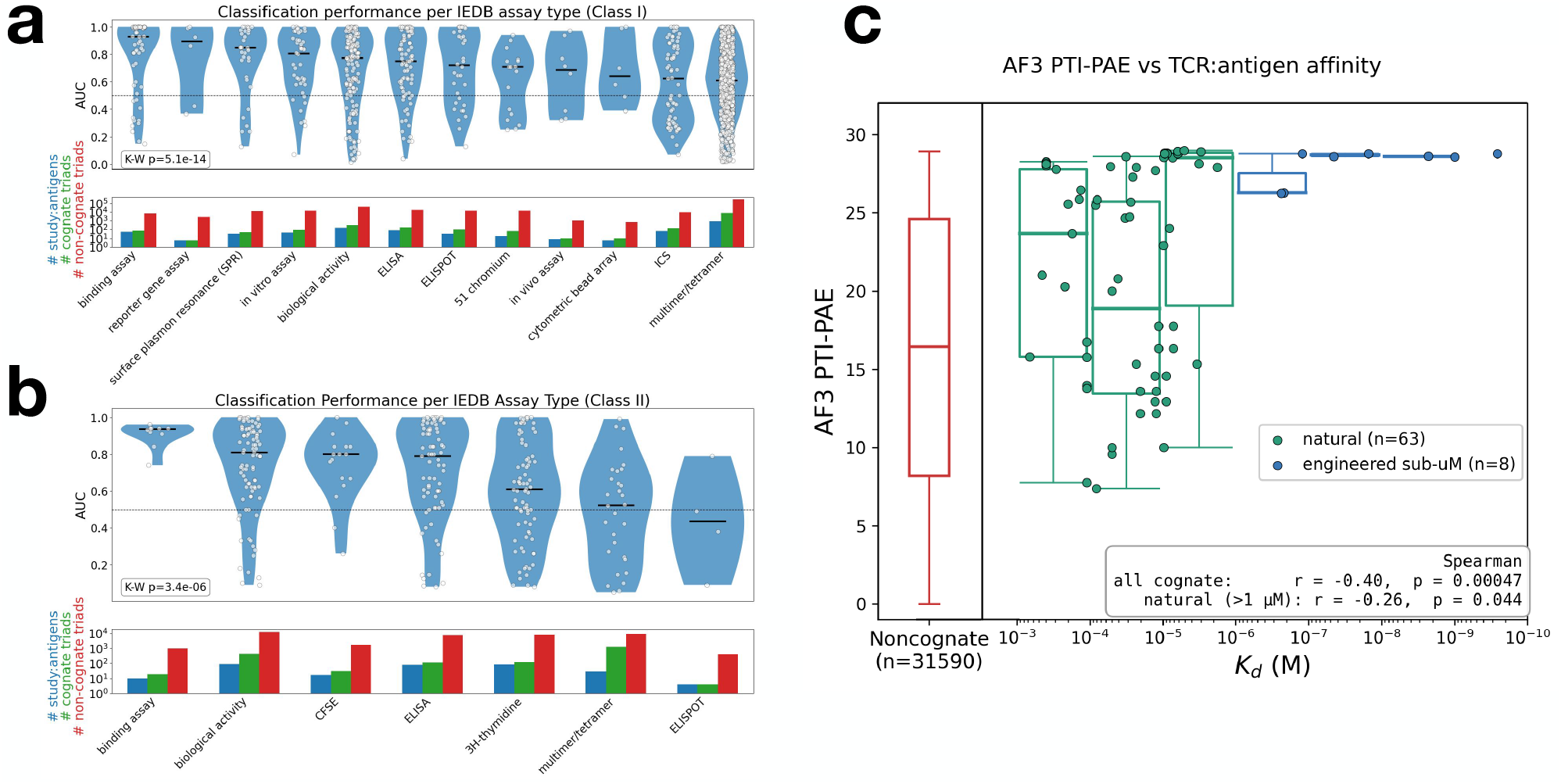
Prediction performance varies by IEDB assay type and TCR:antigen affinity. **(a, b)** The IEDB cognate MHC:peptide:TCR triads described in Figure 3 were classified according to the assay types by which they were identified. Shown, for each assay type containing >3 antigen-study combinations, is the distribution of AUCs (one dot per antigen-study combination) for the respective assay type, calculated using the AF3-PTI-PAE feature (selected based on the Figure 3 analysis). The numbers of antigen-study combinations, and of their constituent cognate and non-cognate triads, for each assay type are shown at the bottom of each plot. **(c)** Relationship between TCR:antigen affinities (*x-axis*) and AF3-PTI-PAE scores (*y-axis*) for cognate triads for which Kds are available in IEDB (n=63 triads with natural TCRs in green, n=8 triads with engineered TCRs in blue), as well as matched non-cognate controls (n=12,350 triads in red). Boxes show AF3-PTI-PAE distributions for each log10 interval of Kd. Statistical comparisons were made for all cognate triads, or all cognate triads with natural TCRs, using Spearman rank correlation test.

The highest-performing assay type for both class I and II was “binding assay” (which comprises cases in which individual TCRs are selected, expressed and assayed for binding) with median AUCs of 0.93 and 0.94, respectively. Also prominent for high performance was Surface Plasmon Resonance (AUC=0.85 for class I). Given their stringent resolution, we expect these assay types to be among the highest confidence approaches for identifying / validating cognate MHC:peptide:TCR triads. Conversely, “multimer/tetramer” – which consists of cellular assays that can have significant background binding when used to identify epitope-specific T cells – was among the poorest performing assay types (median AUCs = 0.61 and 0.52 for classes I and II, respectively), yet contributed the largest number of cognate triads in each case (>80% of class I and >60% of class II). Overall, these observations uncover heterogeneity in the IEDB dataset that could be explained if TCRs identified using different assay types have different degrees of (i) reliability and/or (ii) affinity for their cognate antigens.

To study how the AF3-PTI-PAE score relates to the affinity of the binding interaction, we identified cognate triads in our IEDB set for which experimentally-measured TCR:antigen Kds are available (**Figure 4c**). This consisted of n=67 class I triads and 4 class II triads, with Kds spanning >6 orders of magnitude (range = 260pM–670uM, median = 13uM). Overall, we observed a highly-significant global correlation between AF3-PTI-PAE scores and Kd (p <0.0005) in the expected direction. Particularly notable was a subset of 8 triads with engineered TCRs that had sub-micromolar affinity and whose median AF3-PTI-PAE score ranked at the 6th %ile of the 63 natural TCR Kds analyzed. Excluding this subset, and focusing exclusively on natural TCRs, the correlation between AF3-PTI-PAE and Kd remained significant (p=0.044). These results indicate that the performance of the classification model is dependent on affinity within the physiological range, and that it appears to achieve even higher accuracy for affinity-matured TCRs.

We also performed an exploratory assessment of the AF3-PTI-PAE model’s performance on diverse peptides that are cross-reactive for a fixed TCR. Within our IEDB dataset, we focused on the TCR for which the largest number of different cognate peptides had been identified. This corresponded to the “AGA1” TCR recognizing the HIV Gag peptide KAFSPEVIPMF in complex with the B*57 allele (associated with natural control of infection), that was the basis of a 2020 study by Mendoza et al that used yeast display against a random peptide library to identify diverse cross-reactive peptides [43]. We used AF3-PTI-PAE to score triads containing the following: (i) the original HIV peptide (which is present in PDB structure 2YPL [44]), (ii) 21 cross-reactive peptides identified by yeast display and that were present in our IEDB set (because their sequences were sufficiently far from the original HIV peptide to pass our “seen antigen” homology exclusion filter; range of overall Hamming distances from the original 11mer = 5-9), and (iii) 4,559 background controls in which different numbers of mutations were randomly introduced into the original HIV peptide (range of overall Hamming distances from the original 11mer = 1-11).

While many minimally-mutated peptides showed signal approaching that of the original peptide (potentially attributable to a combination of both true cross-reactivity and false positive scoring by AF3-PTI-PAE), this signal progressively waned as the distance from the original peptide increased **(Supplementary Figure 2, *pink v green*)**. However, when matched for mutational distance, the experimentally-defined cross-reactive peptides scored much more confidently than the corresponding randomized peptides **(Supplementary Figure 2, *blue v pink*)**. This illustrates that the model can successfully recognize diverse cross-reactive peptides for a fixed TCR against the null hypothesis that all mutations affect binding equally.

### Classifier validation on high-quality held-out datasets

To estimate the generalizability of AF3-based classifier, we turned to our held-out validation set. For class I, this consisted of cognate triads from the PDB that were released after the AF3 training cutoff date (January 12, 2023) and identified in TCR3d. This included a total of 16 cognate triads (representing 7 unique MHCs and 14 unique p:MHC antigens), together with 181 permuted non-cognate controls. Of the 14 unique p:MHC antigens, 6 had high-homology matches within the PDB prior to AF3 training, however when adding the TCR, none had substantial overall homology to a triad in the pre-AF3 PDB dataset (per our exclusion criteria described above).

For class II, the validation set consisted of triads that we recently identified with an assay (the Clonally Resolved Epitope and Transcriptome Assay – CRESTA) that uses multiplexed DNA-barcoded p:MHC probes and single cell sequencing of clonally expanded cells to map TCRs across many class II-restricted epitopes simultaneously(^45^). Importantly, unlike typical single cell approaches, CRESTA uses concordant probe signals from ≥3 individual cells all sharing the same TCRα:β pair, resulting in high-confidence TCR:epitope assignments. We used the 205 previously-described CRESTA cognates triads, together with 1228 matched non-cognate controls. These triads are distributed across a total of 8 MHC class II-restricted antigens from *Mycobacterium tuberculosis*, none of which has been structurally characterized (hence all 8 are strictly “unseen antigens”).

We applied the AF3-PTI-PAE model to triads in these validation sets, and performed ROC analysis for each unique antigen (14 antigens for class I, 8 antigens for class II) (**Figure 5a,b**). This analysis yielded AUC values of 0.50-1.00 (median 0.92) for class I, and 0.71-0.91 (median 0.81) for class II, representing unprecedented performance for this challenging problem. We also observed that a subset of cognate triads for each class was detectable at very high specificity: 5 of 16 (31%) class I cognate triads were ranked above all (≥12) non-cognate controls for their respective antigens, and for class II the model achieved median sensitivities across the 8 antigens of 13%, 26% and 52%, respectively, at 100%, 99% and 95% specificity. Precision-recall analysis yielded median average precisions of 0.50 and 0.56 for class I and II, respectively (**Supplementary Figure 3a**).

**Figure 5:**
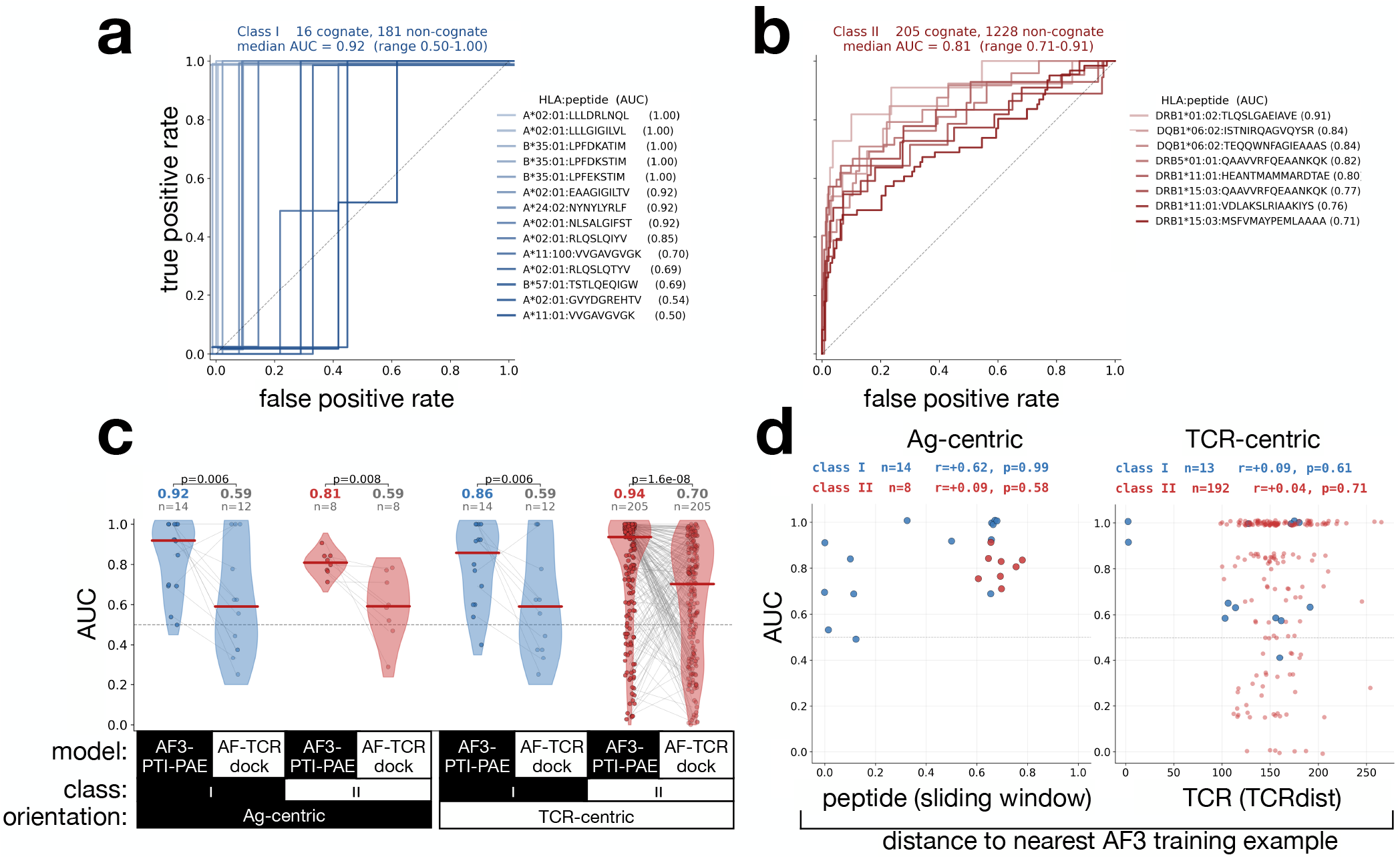
Validation of AF3-PTI-PAE classification performance on held-out datasets of high-quality MHC I and II triads. **a, b** ROC analysis applying the best model developed in Figure 3 (AF3-PTI-PAE) to sets of high-quality, held-out triads (all released after the AF3 training cutoff date): 14 MHC class I antigens with 16 cognate and 181 non-cognate triads from the PDB, and 8 MHC class II antigens with 205 cognate and 1228 non-cognate triads from CRESTA^45^. Lines show the classification performance across cognate v non-cognate triads for each antigen individually. Values outside of the 0–1 interval in (a) result from the use of jitter to more easily visualize overlapping ROCs. **(c)** Violin plots comparing the classification performance (AUC) of AF3-PTI-PAE v state-of-the-art (AF-TCRdock – [31]) on the class I and II validation triads described in (a, b). Each model was evaluated in the antigen-centric orientation described in (a, b), as well as a TCR-centric orientation in which cognate v non-cognate antigens were compared for each TCR. Lines connect corresponding datapoints between the 2 models, and p-values were calculated using the Wilcoxon signed-rank test. Two class I values could not be calculated for AF-TCRdock because they involved mixed species (mouse TCRs specific for human MHCs). **(d)** To assess the generality of classification performance, we compared each of the AF3-PTI-PAE-based AUCs described in (c) against distance to the nearest AF3 training example. For antigen-centric AUCs, we use distance to the nearest training peptide measured by sliding window identity (left), and for TCR-centric AUCs, we use distance to the nearest training TCR measured by TCRdist (*right*). P-values were calculated using the Spearman rank correlation test.

In addition to these antigen-centric scores, in a secondary analysis we assessed performance of the AF3-PTI-PAE model on the converse (TCR-centric) formulation of the problem, reflecting another important use-case of these models. Using the same validation dataset, we built ROCs for each TCR, this time comparing the scores of cognate v non-cognate antigens. Here again we observed high performance, with median AUCs of 0.86 and 0.94, for classes I and II, respectively (**Figure 5c**). Importantly, for both the antigen- and TCR-centric analyses, we found that AF3-PTI-PAE substantially outperformed (by 0.24-0.33 AUC units) the leading AF2-multimer-based model (AF-TCRdock– [31]) (**Figure 5c**). The secondary models explored earlier (Random Forest and XGBoost, in **Supplementary Figure 1c**) also outperformed AF-TCRdock, but were weaker than AF3-PTI-PAE, particularly for class II where we had suspected overfitting (median AUCs of 0.83 and 0.63, for class I and II, respectively – **Supplementary Figure 3b**).

Finally, to stringently assess the generality of the model beyond its training data, we tested whether performance (measured by antigen- or TCR-centric AUCs) correlated with distance between our validation triads and the pre-AF3 PDB data (measured by peptide sequence distance for the antigen-centric analysis, or the TCRdist [18] measure of CDR homology for the TCR-centric analysis). In both formulations, and across both MHC classes, we observed no correlation between performance and distance to AF3 training data (**Figure 5d**). For the class I PDB triads we also found that the AF3-PTI-PAE metric was correlated with structural accuracy (measured by RMSD relative to the ground truth crystal structures – (**Supplementary Figure 3c**)), consistent with the interpretation that the model’s performance is attributable to accurate structural prediction of cognate triads.

## Discussion

In this study, we show that AF3 can model the MHC:peptide:TCR complex with sufficient accuracy and generality to enable a solution to the problem of predicting TCR specificity against “unseen” antigens from sequence alone. We begin with a benchmarking analysis on PDB data, in which we find that AF3 can reproduce the ternary structures of known MHC:peptide:TCR triads, including triads on which it was not trained, with state-of-the-art accuracy (**Figure 1**). We find that this performance is strongly correlated with model error metrics, supporting the use of such metrics as features for the prediction of specificity on triads of unknown structure. Next, by applying AF3 to a much larger and more diverse set of triads lacking prior structural characterization, and comprehensively screening 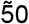 features of the predicted structures, we construct a general model for distinguishing cognate from non-cognate TCRs (**Figure 3**). We find that this model has unprecedented performance on independent high-quality datasets for both MHC classes (**Figure 5**).

Given the relative simplicity of our final AF3-PTI-PAE model, and its stability during validation, we expect its performance to generalize well to other cases, and to be immediately useful for querying TCRs of interest against small-to-moderate-sized sets of candidate epitopes. However, while these results represent a major advance over the essentially random performance that is the current state-of-the-art^37,46^, the current model is unlikely to be well-suited to applications requiring high sensitivity or involving very large repertoires in which the prior probability of any given TCR being specific for a query antigen is very low. Nonetheless, it is notable that we frequently detect a subset of cognate TCRs at high specificity– for example, a median of 13% sensitivity at 100% specificity, and 83% sensitivity at 50% specificity for our class II validation set (**Figure 5**) – with the last ~50% of cognate triads distributed near-randomly among the non-cognates. This suggests that AF3-PTI-PAE may nonetheless be useful for some repertoire-scale applications that can tolerate low individual-epitope sensitivity (e.g. because multiple epitopes from a protein / organism are being integrated together). It also indicates that the model’s overall performance is likely limited by a failure of AF3 to accurately predict the structures of a subset of cognate triads.

More generally, the classification performance that we demonstrate here serves as a baseline above which we expect improvements will be made via at least two mechanisms. First, as general structural prediction models continue to evolve, and especially as their modeling of multi-protein complexes becomes more accurate, we expect these enhancements to translate directly to improved performance for the TCR specificity problem. Second, and perhaps more powerfully, there is potential for fine-tuning / adapting of these general structural prediction approaches to the particular task of TCR specificity prediction, for example by fine-tuning on triads with solved structures using an auxiliary loss that penalizes deviations between predicted and experimental structures (e.g., RMSD). An alternative strategy requiring substantially fewer trainable parameters would be to retain AlphaFold3 as a frozen structure generator and instead learn a lightweight *post hoc* correction that predicts rigid-body transformations applied to either the TCR, the pMHC complex, or both. In this framework, AF3 is used to generate high-quality individual structures for the TCR and the peptide:MHC complex (supported by our results in **Figure 1c**) after which a regression model predicts optimal rigid transformations (rotations and translations) to refine their relative orientation.

The success of fine-tuning models to increase performance on the general MHC:peptide:TCR problem, however, will depend heavily on the availability of sufficiently large and accurate datasets of confidently-resolved MHC:peptide:TCRαβ triads. In this regard, our finding that the AF3 PAE-PTI model’s performance is heterogeneous across IEDB studies and assay types (**Figure 4**) is especially notable, and builds on a recent observation that the expected reactivities could not be reproduced in ~50% of TCRs in a similar database^42,47^

In addition to the potential for inaccuracies within TCR databases, a second, non-mutually exclusive explanation for the assay-to-assay differences that we observe in **Figure 4a,b** could be that the AF3 PAE-PTI model is differentially sensitive across the affinity range, and that different assay types detect (and/or are applied to) triads of different TCR:antigen affinities. We find support for the first of these propositions in our analysis of IEDB triads for which binding affinity measurements are available, where we observe significant variation in the AF3-PTI-PAE score across the physiological range of TCR affinities (**Figure 4c**). Interestingly, we also consistently observe especially strong AF3-PTI-PAE scores for engineered TCRs that have nanomolar affinity, suggesting that the relatively low (micromolar) affinity of physiological TCRs for their antigen ligands may impose a fundamental limit on the sensitivity of the approach, relative to cases like the antibody-antigen interaction that often involve higher affinities. On the other hand, the contribution of the MHC, which fixes the linear conformation of peptide and has less overall structural diversity than an antibody epitope, may help to confine the effective search space relative to the antibody-antigen problem.

Despite these advances, the most challenging problem – accurate, unconstrained *in silico* mapping of a query TCR – remains out of reach, insofar as it requires a false-positive rate that is commensurate with the billions of p:MHC antigen possibilities, and is ultimately limited by the poly-reactivity of individual TCRs for diverse peptides^48^. Nonetheless, it is conceivable that the improvements described above, potentially in combination with other approaches (such as protein language models[49]), could one day approach such performance. It is also possible that generative approaches such as the RFdiffusion models^50–52^ could in future be applied in a way that replaces billions of antigen-specific queries with the single design problem of generating a p:MHC (under certain constraints) to bind the TCR of interest, and then using it to identify plausible candidates based on homology or other criteria. Finally, while our focus here has been peptide epitopes restricted by classical class I and II MHCs and recognized by alphabeta TCRs, the generality of AF3 suggests that the same type of approach may be extensible to non-peptidic ligands, nonclassical MHCs and/or other classes of TCRs; and more generally, to other, non-immune hypervariable molecular interfaces.

## Methods

### AlphaFold 3 configuration

AlphaFold 3 was run with MSAs and templates turned on. No custom templates or alignments were used. All triads were run with 5 seeds (1, 2, 3, 4, 5) and 1 diffusion sample due to the observation that increasing the number of seeds results in greater model accuracy improvements than increasing the number of diffusion samples^38^. The default number of recycles was used (10).

### Boltz-2 configuration

Boltz-2 was run with MSAs turned on and templates turned off (default behavior). The number of recycling steps was increased to 5 (from the default 3) and diffusion samples was increased to 5 (from the default 1) as was done in the Boltz-2 PDB baseline^40^.

### RMSD measures

*RMSD* is defined as the root-mean-square deviation between two corresponding sets of atom positions after optimal rigid-body superposition.

*MHC RMSD* is defined as the protein backbone RMSD of the MHC *α* and *β* chain for MHC class II and RMSD of the MHC *α* chain for MHC class I (with the reasoning that *β*_2_-micro-globulin is not involved in the binding site) after superposition of the chains. In some cases, the MHC contains a flexible linker sequence to the peptide. This sequence is not included when computing MHC RMSD.

TCR RMSD is defined as the protein backbone RMSD of the IMGT-numbered V-region of the TCR chains. IMGT numbering is performed using ANARCI^53^. Only this region is considered because the C-region is distant from the binding site.

*Peptide RMSD* is defined as the RMSD of the peptide alpha carbons between the predicted and crystal structures after aligning the MHC coordinate frames (which are extracted using TCRdock^31^). In contrast to earlier work^31^, we use alpha carbons instead of a full backbone RMSD to compute the metric consistently even for analyzed PDB entries which have missing backbone residues in their peptides.

*CDR RMSD* is defined as RMSD of the predicted and crystal CDR regions of the TCRs after alignment of the MHC coordinate frames, with the CDR3 region upweighted by a factor of 3^31^. IMGT numbering via ANARCI^53^ is applied to identify the CDR regions. Although exact IMGT definitions of the CDR regions differ in the literature, the calculation for CDR RMSD in this paper used the regions defined as the inclusive ranges of IMGT-numbered residues (27, 38), (56, 65), (81, 86), and (104, 118) for the CDR1, CDR2, CDR2.5, and CDR3 regions, respectively.

In all computed RMSD values, small discrepancies between the residue sequence in the PDB structure file and the full fasta sequence for the PDB entry (which is used for prediction) are accounted for using Biopython’s `Bio.pairwise2.align` module^54^ to sequence align predicted and crystal chains before computing RMSD.

### Data exclusions

To maximize generality, triads were excluded from the training set if they had a substantial overlap with either (i) pre-AF3-cutoff PDB triads (to avoid cases where AF3 had been trained on the structure of the same or a similar triad) or (ii) their respective MHC class validation sets (to preserve the separation between our own training and validation sets). “Substantial overlap” here means having sequence identity match of ≥ 95% for both MHC and both TCR chains and sequence identity match of ≥ 77% for the peptides (e.g. corresponding to ≥7 identical amino acids for a 9mer peptide). Additionally, we removed one additional triad from the class I training set which had a post-AF3-training sequence identity match but which was not queried by STCRdab.

### Design of non-cognate triads by permutation

Given a set of cognate triads that share the same antigen in a dataset, we generated an excess of non-cognate triads via a permutation approach using other triads in the same dataset. For each antigen, we sought to design at least a 10-fold excess of non-cognate triads, and increased this ratio even further to preserve resolution for antigens with relatively few binding TCRs (specifically, for a p:MHC complex with *n* cognate TCRs, we generated max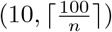 non-cognate triads). In rare cases where the number of cognate triads was very large and the target number of non-cognates was not possible within the dataset, we generated the maximum possible number of non-cognate triads (also allowing for the exclusion criteria, described below).

Our approach assumes that the probability of a TCR being specific for more than one antigen within the database is generally very low. To further control this possibility (by removing permuted p:MHC and TCR pairings at higher risk of being accidental cognates), we designed a set of exclusion criteria that we applied to permuted non-cognates. After first permuting all possible pairs, we kept only the non-cognate pairings in which:

1. The permuted TCR’s original (known binding) p:MHC complex has a total (MHC sequence and peptide sequence) edit distance of at least 3 from the permuted p:MHC complex.
2. The permuted TCR, and all TCRs that are known to bind the same p:MHC complex, have a TCRdist^18^ of at least 120 from all the TCRs known to bind the permuted p:MHC complex.

After generating all possible negatives under these rules, we sampled down to the target number of cognate triads per each p:MHC complex by first randomly selecting an “antigen of origin” and then randomly sampling a TCR from that antigen in order to prevent over-sampling TCRs from over-represented antigens in each dataset. To prevent duplicate non-cognate triads, a TCR is only sample-able once per p:MHC complex, even if multiple antigens-of-origin contain it.

### Docking geometry Z-score and probability

Z-score of docking geometry is calculated by first creating a consensus distribution of docking geometries from the PDB triad geometry databases in the TCRdock repository^31^. Each 6-parameter docking geometry (distance, torsion, TCR unit y, TCR unit z, MHC unit y, and MHC unit z) is first converted into a 7-parameter docking geometry by converting torsion from radians into a sin-cos encoding so that torsion values close to 0 and close to 2*π* are encoded similarly. Then, for each database, the mean and covariance matrix of the seven features are estimated to define the consensus distribution.

Next, given a true or predicted triad structure and its 6-parameter docking geometry, a “Z-score” can be taken by similarly converting to 7 parameters and then taking the Mahalanobis distance from the consensus distribution for the MHC class of the triad. The Mahalanobis distance can be converted into a p-value using the formula:

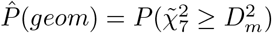

Where 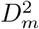 is the squared Mahalanobis distance.

### DOCKING metrics

Docking metrics (d, torsion, tcr_unit_y, tcr_unit_z, mhc_unit_y, mhc_unit_z) are used unmodified from TCR-dock output^31^. “pred_dgeom” is described above.

### Local metrics

Local metrics (ipTM, ranking score, and pTM) are used unmodified from AF3 output^38^.

### Interface metrics

Interface metrics are named using the following nomenclature:

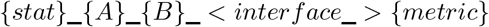

Where *stat* is either “min” or “mean”. *A* and *B* are regions of the metric matrix corresponding to either the peptide (“p”), the MHC chains (“mhc”), the MHC and peptide chains (“pmhc”), or the TCR chains (“tcr”). “mhc” refers to the *α* and *β* residues for class II or only MHC *α* residues for class I. “pmhc” refers to the same region as “mhc” but with peptide residues as well. “tcr” refers to the the V-regions of the TCR chains as identified by ANARCI IMGT numbering^53^. Whenever the “interface” keyword is present, non-peptide regions are collapsed to only the residues within 8 angstroms of the peptide.

*metric* refers to either the PAE or contact probability matrix. Note that the PAE matrix is not symmetric (unlike the contact probability matrix), so “mean_tcr_p_pae” is not equivalent to “mean_p_tcr_pae”.

Two exceptions exist to this naming rule:

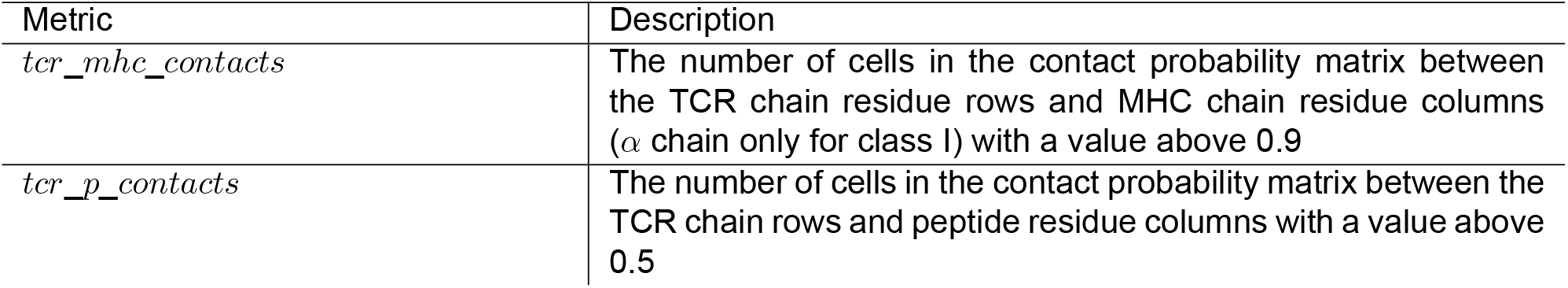

### Local metrics

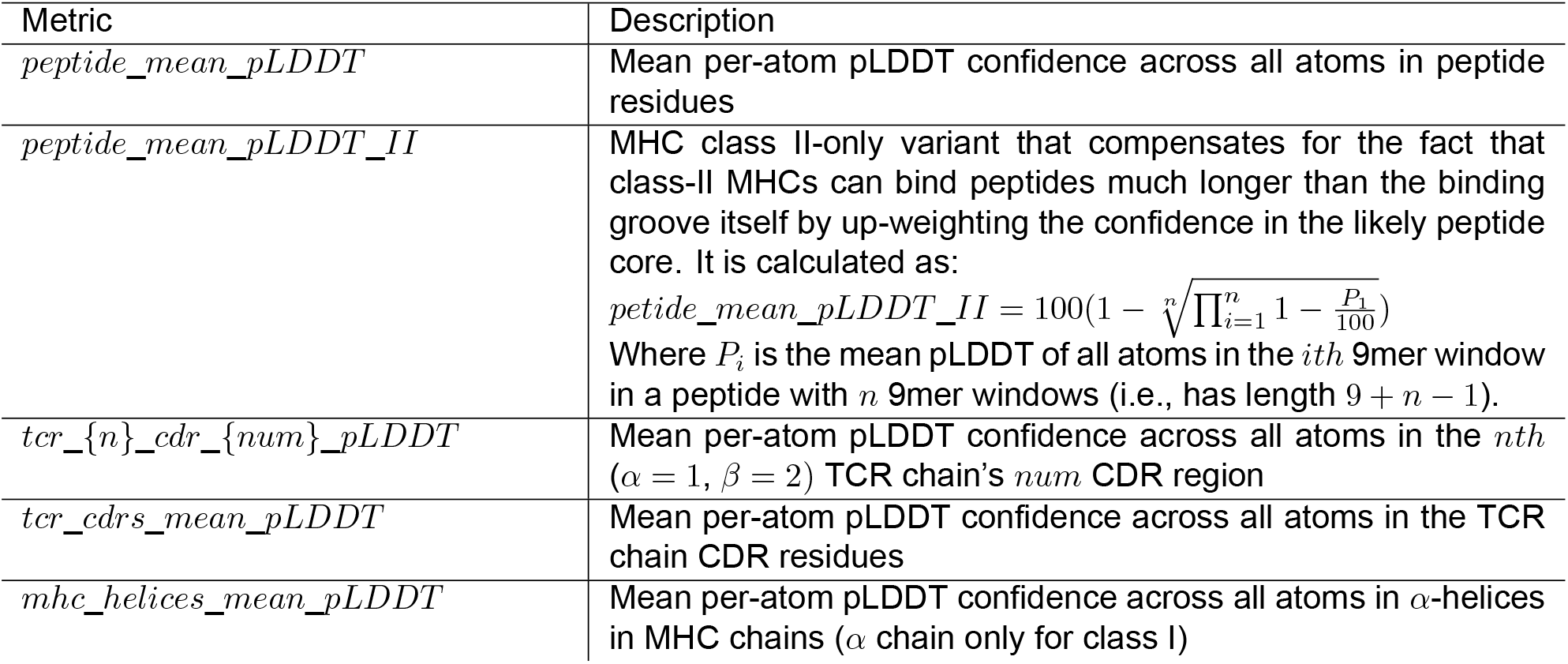

### Machine learning models

Random Forest and XGBoost models were developed with 5-fold cross-validation. To minimize information leakage, folds were defined at the level of antigen-study combinations, such that all cognate and non-cognate triads associated with a given antigen-study combination were assigned to the same fold. Performance was evaluated using the same antigen-study-combination-level ROC/AUC framework used throughout the manuscript. For each MHC class, we compared 11 candidate models in total: the original PTI-PAE classifier, together with Random Forest and XGBoost classifiers trained using (i) all AF3-derived features,(ii) summary features only, (iii) interface features only, (iv) local features only, and (v) docking features only.

## Software and data availability

All analysis code and original data (excluding inference output) for the paper is available at https://github.com/AltinLab/tcrtrifold-experiments.

## Acknowledgments

We are grateful to Joel Ernst for helpful discussions about this work. This work was supported by the following NIH awards: R21AI149311, R01AI173002, 75N93024C00054.

## Supplementary Figures

**Figure S1:**
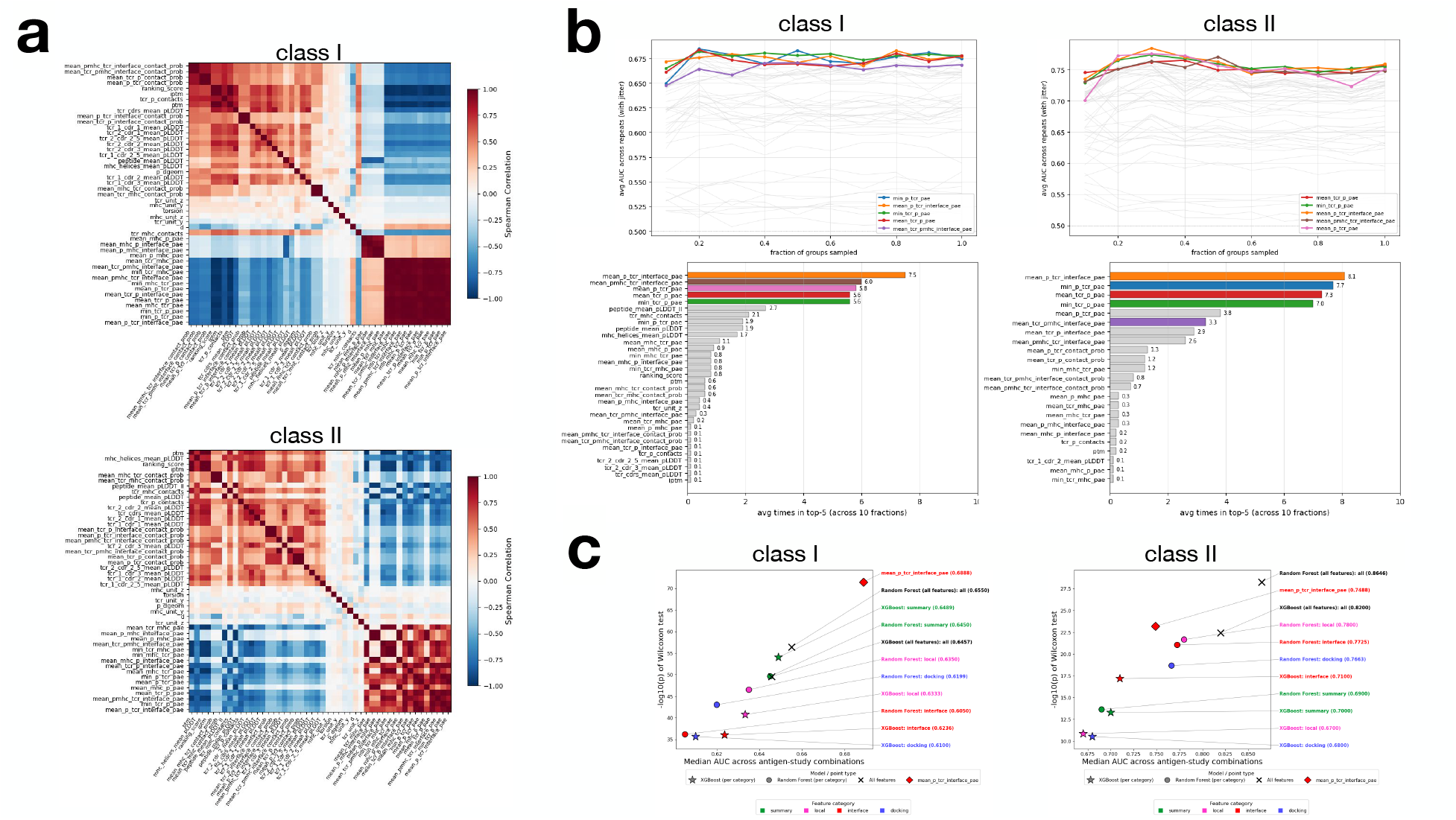
Stability of individual feature performance within the IEDB training data, and comparison against alternative machine learning models. **(a)**Feature-feature correlation analysis for the IEDB dataset shown in Figure 3, calculated based on Spearman comparisons. **(b)** Stability of the features described in Figure 3 across randomized subsamples of the IEDB dataset at increasing fractions of the available data (upper plots, x-axis). At each fraction, we generated 10 independent random subsets, ranked all features according to their median antigen-study-combination AUC, and then averaged the results across runs (lower plots). **(c)** Performance of Random Forest and XGBoost classifiers trained on the IEDB dataset. Models used either all features within a particular class (denoted using the color scheme of Figure 3), or all features within all classes (black). Models were trained and evaluated using 5-fold cross-validation with folds defined such that all cognate and non-cognate triads associated with a given antigen-study combination were assigned to the same fold. Performance was evaluated using the same antigen-study-combination ROC framework used in Figure 3. In all panels, the feature selected for downstream analysis (AF3-PTI-PAE) is named “*mean_p_tcr_interface_pae*”; it was added to panel (c) as a point of comparison (red diamond).

**Figure S2:**
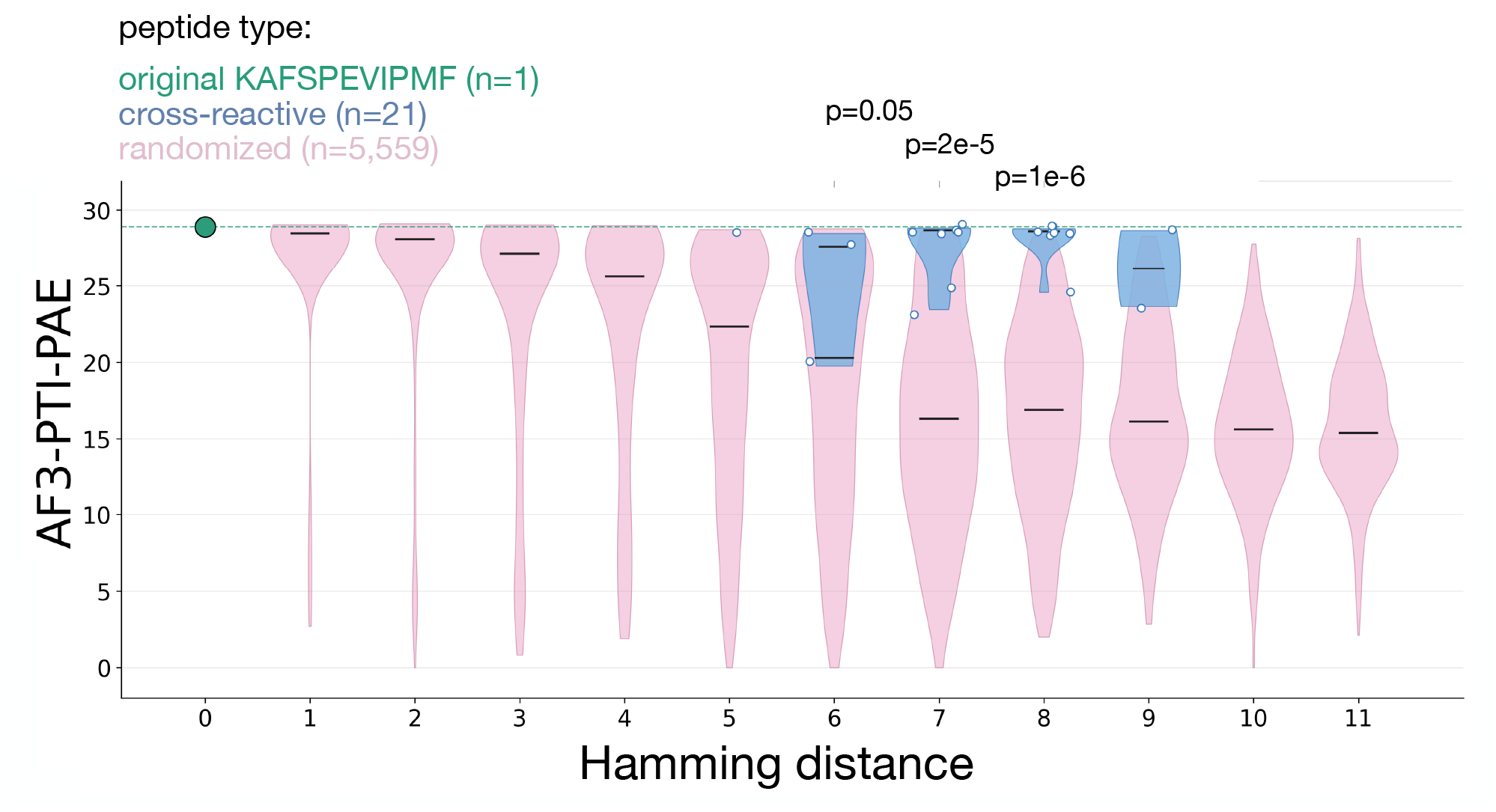
AF3-PTI-PAE analysis of peptide cross-reactivity to a fixed TCR. A set a 21 peptides, identified previously using yeast display [43] as being recognized in the context of HLA-B*27:03 by the HIV-specific TCR “AGA1”, were the largest set of cross-reactive antigens for a single TCR in our IEDB dataset. Shown for the original HIV peptide (green), the 21 cross-reactive peptides (blue), and 4,559 controls randomly-mutated to a range of distances (pink), are the AF3-PTI-PAE scores for the corresponding triads (*y-axis*), plotted as a function of Hamming distance from the original peptide (*x-axis*). For the 3 bins containing ≥3 cross-reactive peptides, p-values were calculated using Wilcoxon Rank-Sum tests.

**Figure S3:**
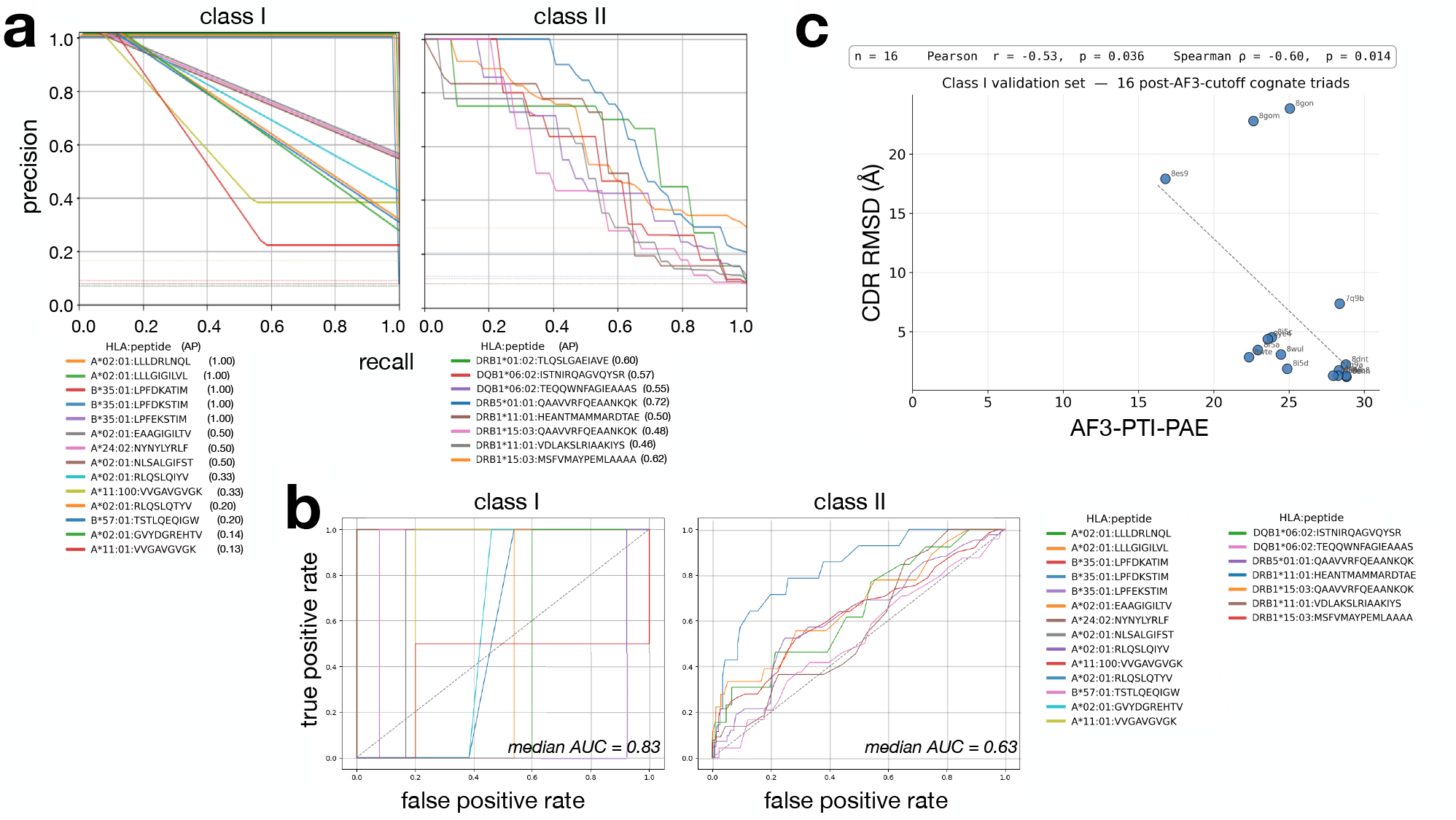
Further analysis of the validation dataset from Figure 5. **(a)** Antigen-centric Precision-Recall analysis of the class I and II validation triads. AP denotes Average Precision. **(b)** Antigen-centric ROC curves showing the performance of the *Random Forest (all features)* model described in Supplementary Figure 1c, on the validation dataset. **(c)** Correlation between AF3-PTI-PAE model score and structural accuracy (CDR RMSD) for all validation triads with available crystal structures. PDB identifiers are shown next to each datapoint.

## Notes

### Competing Interest Statement

The authors have declared no competing interest.

